# Reversible disulfide bond crosslinks as tunable levers of phase separation in designer biomolecular condensates

**DOI:** 10.1101/2024.07.13.603402

**Authors:** Malay Mondal, Penelope E. Jankoski, Landon D. Lee, Daniel M. Dinakarapandian, Tzu-Ying Chiu, Windfield S. Swetman, Hongwei Wu, Anant K. Paravastu, Tristan D. Clemons, Vijayaraghavan Rangachari

## Abstract

Biomolecular condensates (BCs) are membraneless hubs enriched in proteins and nucleic acids that have become important players in many cellular functions. Uncovering the sequence determinants of proteins for phase separation is important in understanding the biophysical and biochemical properties of BCs. Despite significant discoveries in the last decade, the role of cysteine residues in BC formation and dissolution has remained unknown. Here, to determine the involvement of disulfide crosslinks and their redox sensitivity in BCs, we designed a ‘stickers and spacers’ model of phase-separating peptides interspersed with cysteines. Through biophysical investigations, we learned that cysteines promote liquid-liquid phase separation in oxidizing conditions and perpetuate liquid condensates through disulfide crosslinks, which can be reversibly tuned with redox chemistry. By varying the composition of cysteines, subtle but distinct changes in the viscoelastic behavior of the condensates were observed. Empirically, we conclude that cysteines are neither stickers nor spacers but function as covalent nodes to lower the effective concentrations for sticker interactions and inhibit system-spanning percolation networks. Together, we unmask the role of cysteines in protein phase behavior and the potential to develop tunable, redox-sensitive viscoelastic materials.

## INTRODUCTION

Biomolecular condensates (BCs) are dense hubs of membraneless organelles commonly containing proteins and nucleic acids that are ubiquitously observed in cells across all kingdoms of life and are speculated to have been present as protocells during the early origins of life on Earth ^1–6^. BCs reversibly achieve need-based spatiotemporal organization and control of cellular matter in an energy-independent manner^7–10^. BC formation is a density transition in which a denser phase (or phases) enriched in biomolecules coexists with a biomolecule-deplete dilute phase above a threshold saturation concentration (*C_sat_*)^11^. The coacervation of biomolecules toward such a density transition is better captured by a phenomenon called liquid-liquid phase separation (LLPS). Two types of coacervation predominate BCs. Self-coacervation is unimolecular, involving polypeptides undergoing LLPS by themselves in specific ionic strengths, while complex-coacervation involves biomolecular scaffolds that partition other interacting partners called clients within the condensates^12^.

Irrespective of the coacervation type, the BCs’ interactions involve weak, non-stoichiometric multivalent interactions, including cation-π, π-π, Van der Waals, and hydrogen bonding.^12–14^ LLPS among associate polymers such as proteins, is best defined by a ‘stickers and spacers’ model in which stickers are amino acid residues that are involved in weak, non-covalent interactions with one another, that are interspersed with disordered spacers that contain non-interacting residues^13,15^. The balance of the interaction strength between the stickers and the spacers’ effective solvation volume determines the viscoelastic properties of the BCs formed^16^.

A spectacular array of functions that BCs are known to be involved in has also inspired researchers to develop them into dynamic compartments and soft materials for many biotechnological and pharmaceutical applications^17–23^. Associative biopolymers such as proteins, either by self-coacervation or by complex-coacervation with clients such as RNA, form BCs best explained by the process of LLPS^11,24,25^. Several researchers have recently exploited molecular features of condensates such as intrinsic protein disorder, composition and sticker valence, spacer scaffold, and client chemistry to design tunable viscoelastic materials to cater to a wide range of functionalities^21,22,26–28^. However, understanding the design and properties of redox-sensitive condensates remains limited^19,29,30^. Only a handful of studies have investigated redox sensitivity in peptides and proteins based on the oxidation of methionine to sulfoxides and sulphones^29,31^. The dearth of information is especially apparent in the use of cysteines as redox-modulating residues within the peptide sequence. Cysteines are considered to be order-promoting amino acids due to their ability to form covalent disulfide bonds^32–35^. Therefore, it may seem counter-intuitive to see the presence of cysteines among disorder-promoting amino acids such as arginine, glycine, and serine, etc. However, nature seems to have accommodated disorder-promoting sequences interspersed with cysteine, especially in complex higher-order organisms such as eukaryotes, which are often involved in reactions with redox flux^35^. However, the role of cysteines in the formation and dissolution of BCs has remained unknown. Previously, we demonstrated that cysteine-rich protein modules called granulins (GRNs) modulate LLPS of TAR-DNA binding protein (TDP-43) by tuning the redox state of cysteine^36,37^, suggesting that cysteine could play a role in coacervation and LLPS. Recently, cystamine-linked peptide synthons were used to demonstrate the effectiveness of redox-sensitive phase-separating peptides^28^, which further elucidates the potential of cysteines to be used in designer redox-sensitive biomolecular condensates.

Here, we set out to understand the following important questions: *i*) Do covalent crosslinks formed by disulfide bonds facilitate LLPS? *ii*) Do disulfide bonds function as covalent stickers, spacers, or extended networks? And *iii*) Do BCs formed show sensitivity to redox flux? By designing short peptides containing classical stickers and spacers interspersed with cysteine with varying positional and compositional biases, we learned that cysteine disulfide bonds promote LLPS reversibly, providing intriguing new clues to understanding the dynamics of BC formation and dissolution dynamics.

## METHODS

### Materials

Rink Amide Protide Resin, Fmoc-protected amino acids, and ethyl cyanoglyoxlate-2-oxime (Oxyma) were purchased from CEM peptides. Dichloromethane (DCM), diethyl ether, trifluoroacetic acid (TFA), N-dimethylformamide (DMF), acetonitrile, diisopropylcarbodiimide (DIC), triisopropylsilane (TIS), ethane-1,2-dithiol (EDT), and all other solvents were purchased from ThermoFisher Scientific (USA) or Sigma Aldrich Corporation (USA) at the highest purity.

### Peptide Synthesis

Peptides were synthesized on a Liberty Blue 2.0 automated peptide synthesizer (CEM) through standard 9-fluorenyl methoxycarbonyl (Fmoc)-based solid phase peptide synthesis. Peptide synthesis was performed at a 0.25 mmol scale using Rink Amide Protide Resin (0.65 mmol/g loading, 100-200 mesh). Deprotection of Fmoc protecting groups was carried out using 20 v/v% piperidine in DMF. Each amino acid addition was carried out using Fmoc-protected amino acids (0.2 M), DIC (1M), and Oxyma (1 M) in DMF. After the final Fmoc deprotection, the resin beads were washed 3x using DCM. The peptide underwent global deprotection and cleavage from the resin beads through gentle shaking in TFA/TIS/H_2_O/EDT (92.5: 2.5:2.5:2.5) cleavage cocktail for 4 hours at room temperature. Peptides were then precipitated in cold diethyl ether and chilled for four hours at -20 °C. Following this, samples were centrifuged, and the diethyl ether supernatant was decanted from the resulting peptide pellet. The peptide pellet was then resuspended in diethyl ether and chilled overnight at -20 °C. Centrifugation and decanting of the diethyl ether supernatant were performed again before allowing the peptide pellet to air dry. Crude peptides were purified by reverse-phase high-performance liquid chromatography (HPLC) on a Prodigy HPLC system (CEM) with a water/acetonitrile gradient (containing 0.1% TFA). The mass and identity of the eluting fractions containing the desired PSP peptides were confirmed using electrospray ionization (ESI)-mass spectrometry (MS) on a Thermo Scientific Orbitrap Exploris™ 240.

### Turbidity assay

Turbidity measurements were performed on a BioTek Synergy H1 microplate reader. Samples were allowed to equilibrate at room temperature for approximately ten minutes before each measurement. Phase diagrams were generated using a boundary value of 1.40 A.U at 600 nm. Data processing was done using Origin 8.5. Three independent datasets were collected and averaged for each measurement.

### Coating glass slides and coverslips

Microscopic glass slides and coverslips were cleaned with 70% ethanol by sonicating for 15 minutes. Coverslips and glass slides were then allowed to air dry. Glass slides and coverslips were submerged in a coating solution (20% Tween20) for 30 minutes. To remove the extra coating solution, the glass slides and coverslips were rinsed eight to ten times with Milli-Q water. Glass slides and coverslips were then dried at 37°C overnight. Lens paper was used to wrap dry-coated glass slides and coverslips and stored at room temperature until further use.

### Preparation of RNA

Lyophilized powdered Poly-A RNA was acquired from Sigma and dissolved in RNase-free water. Using a conversion factor of 1 absorbance, or 40 µg/mL, the concentration was calculated. The prepared stock was frozen and stored at -80 °C. Prepared stock aliquots were thawed and utilized immediately for experiments.

### Confocal microscopy and FRAP analysis

A Leica STELLARIS-DMI8 microscope at 63x magnification was used to capture confocal microscopy images of the peptide droplets on coated cavity glass slides. In all reactions, droplets were allowed to settle for a few minutes before imaging. 0.5 % FITC was added to the reaction (after forming droplets) for imaging. The internal dynamics of the self and complex condensates were investigated using fluorescence recovery after photobleaching (FRAP). For five seconds, the liquid droplets were exposed to a laser intensity of approximately 90% to achieve photobleaching. The recovery of fluorescence was then observed for 60 seconds. The kinetics of fluorescence recovery were normalized and plotted against time using Origin 8.5.

### Image Processing and Analysis

Confocal microscopy images were processed and analyzed using a custom pipeline implemented in Fiji (version 1.54f) and R (version 4.1.2) to quantify the size distribution of the phase-separated droplets over time. In ImageJ, the images were pre-processed by applying Huang’s auto-thresholding method^38^ for binarization and by removing noise using a despeckle filter. Morphological operations (erosion and dilation) and the watershed algorithm^39^ were used for droplet segmentation. The segmented droplets were analyzed, excluding those touching the edge of the frame, and their properties (area, mean intensity, perimeter, and shape descriptors) were measured and exported as CSV files. In R, the CSV files from Fiji were processed using custom scripts. The tidy verse,^40^ ggsci, and scales packages were utilized for data organization, visualization, and statistical analysis. The droplet counts and size distributions were summarized for each combination of peptide, time point, and replicates. Boxplots were generated to visualize the droplet size distributions across time points for each peptide. The image analysis pipeline assumed that the phase-separated droplets were spherical and did not account for potential deviations from this shape. Additionally, the segmentation process may have introduced errors for closely spaced or overlapping droplets, leading to potential undercounting or inaccurate size measurements.

### Dye partitioning and encapsulation

Peptide solutions were oxidized and diluted as described above. To each solution, 5 µM of dye was added and incubated for 10 minutes. Standard solutions for each equivalent dye and buffer concentration were prepared, and their UV-vis absorbances were recorded. The peptides were centrifuged at 15,000xg for 10 minutes. Of the 100 µL volume, 40 µL was drawn as the supernatant, and the UV-vis absorbance was measured. The encapsulation efficiency of the dye was calculated using the following equation where *A_sup_* is the absorbance of the supernatant, and *A_tot_* is the total absorbance at 490 and 550 nm for fluorescein and TMR-OMe, respectively.

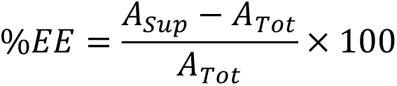

### Circular Dichroism (CD)

CD measurements were performed on a JASCO 810 spectrophotometer. The peptide samples were added to a 0.1 mm path length cuvette and were scanned from 200 to 260 nm. This mixture was allowed to incubate for five minutes before each measurement. Three scans were averaged with a data resolution of 1 nm. The raw data were background subtracted, smoothed with the Slavitky-Golay algorithm, normalized, and plotted using Origin 8.0.

### Nuclear Magnetic Resonance (NMR) Spectroscopy

PSP-1, PSP-2, PSP-3, and PSP-4 peptides were dissolved in 50 mM sodium phosphate (NaPi) buffer and 2.5 mM sodium chloride (NaCl), along with 10% deuterated water (D2O), and pipetting them into 5 mm Wilmad NMR tubes. LLPS samples of all peptides except PSP-5 were first prepared in 3.5 mM peptide concentrations and then diluted to 1.1 mM for non-LLPS samples. The LLPS sample of PSP-5 was prepared in the same buffer at 18 mM concentration, and the non-LLPS sample was obtained by diluting it to 6 mM. The samples also included 10 mM sodium trimethylsilylpropanesulfonate (DSS) as an internal standard for chemical shift calibration. NMR measurements were performed using an 11.75 T magnet (500 MHz, 1H NMR frequency) on a Bruker spectrometer. Using Bruker default “zgpr” pre-saturation pulse sequence, the radio frequency carrier frequency (O1) was optimized to minimize the solvent signal. Then, we used the O1 value on the subsequent excitation sculpting pulse sequence “zgesgp” for water suppression and collected the ^1^H spectra. A recycle delay time (D1) of 1 sec was employed, and signals averaged over 128 scans. All experiments were conducted at a constant temperature of 298K. Spectra were processed by Fourier transform, and ^1^H signal intensities were normalized. Chemical shifts were calibrated for water peak at 4.696 ppm using Topspin 3.7 (Bruker) and analyzed using customized Mathematica scripts.

## RESULTS AND DISCUSSION

### Design of model phase-separating peptides (PSPs)

We set out to answer these questions by designing simple peptide models that recapitulate phase-separating characteristics in associative polymers. First, a control peptide, PSP-1 (Figure 1), was designed based on the stickers and spacers model in which arginines (R) and tyrosines (Y) were used as ‘stickers’ based on their well-known ability to engage in cation-ρε and ρε-ρε interactions in both self- and complex-coacervation modes abundant in BCs^12,13,41–43^. The stickers are interspersed with disorder-promoting glycines (G), and serines (S) called ‘spacers’ that are innocuous to non-covalent interactions but modulate sticker interactions through solvation volume and effective concentrations ^24,25,44^. Using PSP-1 as the basis, cysteine (C) residues were introduced, systematically varying their position and composition (Figure 1). Two cysteines were substituted for serines at the N- and C-terminal ends in PSP-2, while two cysteines replaced two glycine residues in the middle in PSP-3. PSP-4 and PSP-5 were single cysteine variants of PSP-2 and PSP-3 in which the N-terminal cysteine in PSP-2 was retained in PSP-4, and the one in the middle of the spacer from PSP-3 was retained in PSP-5 (Figure 1). Together, the five peptides provided a minimal set of variations to investigate the contributions of cysteines in LLPS. All peptides were synthesized as C-terminal amides (see Methods for details), and target peptides were confirmed by electrospray ionization (ESI) mass spectrometry (Figure S1).

**Figure 1:**
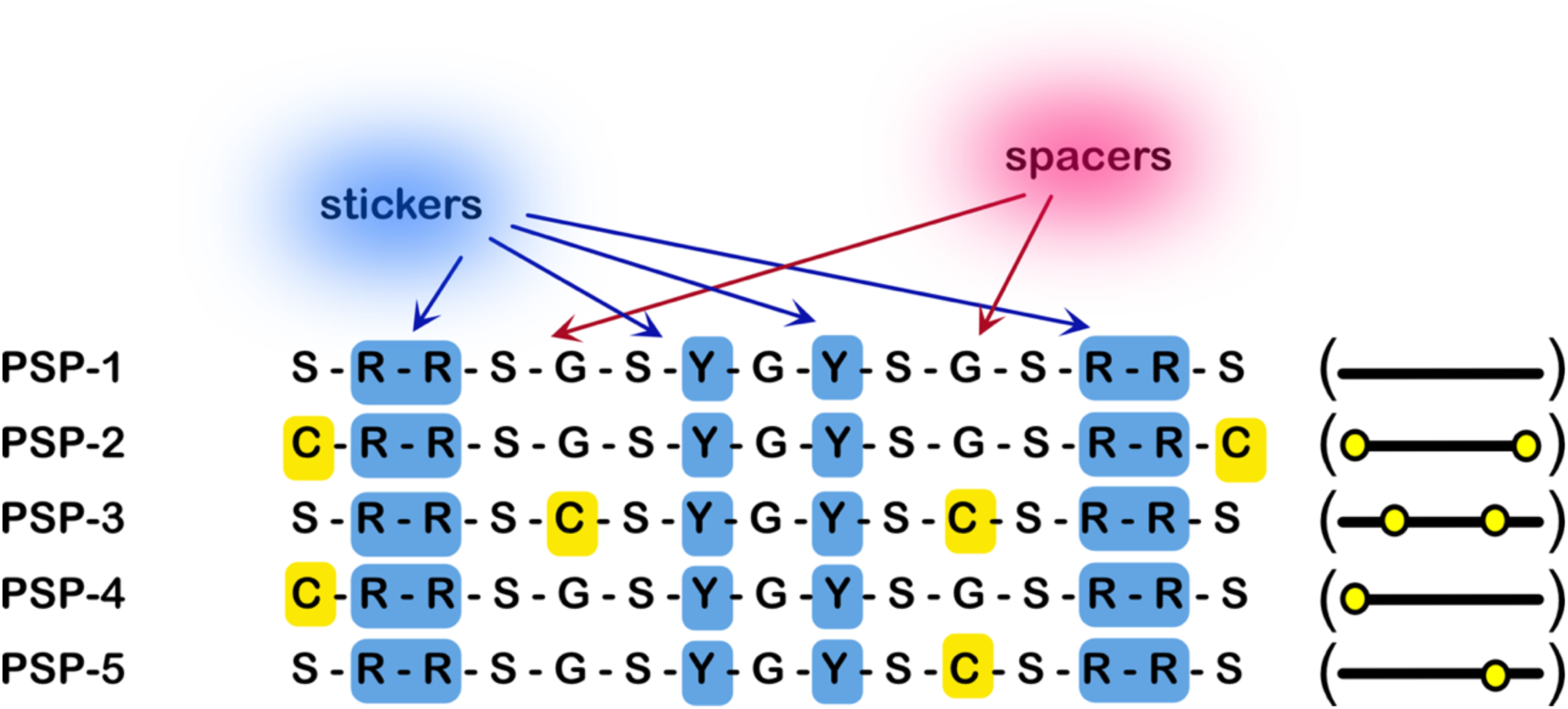
Phase-separating designer peptides used in this study. All peptides were synthesized with a free amine at the N-terminus and an amide at the C-terminus. A schematic representation indicating the positions of the cysteines in the peptide is shown in parenthesis (right).

### Cysteines lower the saturation concentrations for LLPS

The phase transition of a polypeptide from a homogenous to a demixed solution containing two or more coexisting liquid-like phases occurs above a concentration threshold called the saturation concentration (*C_sat_*). We first established phase boundaries and determined apparent *C_sat_* values for the self-coacervates by investigating the concentration, pH, and ionic strength landscape through turbidimetry analysis and confocal microscopy (Figure 2). PSP-1, the control phase-separating peptide without cysteine functionality, was assessed up to 100 mM between pH 7.0 and 12.0 and sodium chloride (NaCl) concentrations of 0 – 3.5 mM (Figures 2a and 2b). The peptide showed LLPS was above 80 mM in 3.5 mM salt concentrations at pH 8.0 upon incubating at room temperature for at least 3 hours and showed a *C_sat_* value of 80 mM in 2.5 mM NaCl (Figure 2b). PSP-2, under similar conditions, showed a dramatic decrease in the phase boundary with LLPS occurring above 2.0 mM and pH 8.0, with an apparent *C_sat_* value of ∼2.0 mM in 2.5 mM NaCl (Figures 2c and 2d). PSP-3 also showed a similar phase boundary in pH scans (Figure 2e) and a slightly altered boundary in ionic strength scans with a *C_sat_* value of ∼1.2 mM in 2.5 mM NaCl (Figure 2f). PSP-4 showed a phase boundary above 3.5 mM and between pH 7 (Figure 2g), with a *C_sat_* value of ∼ 3.2 mM in 2.5 mM NaCl (Figure 2h). PSP-5 demonstrated no evidence of phase separation in the pH scans (Figure 2i) nor in ionic strength scans at or below 3.5 mM peptide concentration (Figure 2j). The peptide showed phase separation only above a *C_sat_* value of ∼16.0 mM in 2.5 mM NaCl with 1% H_2_O_2_ (Figure 2j). The respective differential interference contrast (DIC) and confocal fluorescence microscope images for peptides PSP-1 (Figures 2k and l), PSP-2 (Figures 2m and n), PSP-3 (Figures 2o and p), PSP-4 (Figures 2q and r), and PSP-5 (Figures 2s and t) show the formed droplets along with the turbidity of the phase-separated solution (insets). Fluorescence recovery after photobleaching (FRAP) analysis on PSP-1 droplets showed a somewhat diminished 60% recovery, reflecting less fluidity of the droplets. Droplet fluidity of other peptide coacervates analyzed by FRAP showed ∼80% recovery for PSP-2-5, more fluidity than PSP-1 (Figures 2u-y). Together, these results indicate that cysteines could play an important role in promoting condensate formation, as evidenced by decreased *C_sat_* values of the peptides to phase separate by self-coacervation.

**Figure 2:**
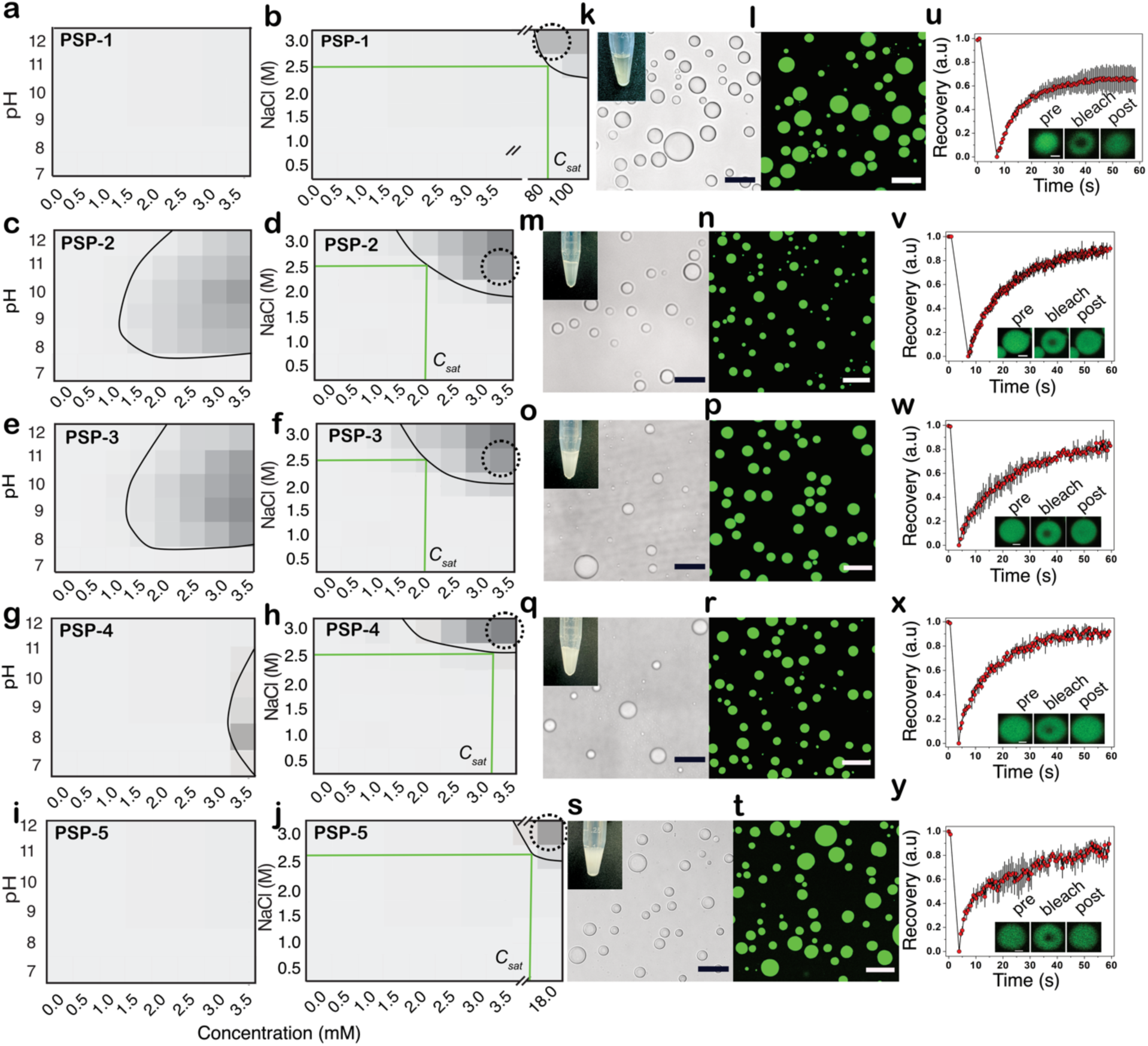
Phase boundaries for peptide self-coacervates. Respective phase diagrams for concentration vs. pH (in 2.5 mM NaCl) and concentration vs. salt NaCl (at pH 8.0) measured based on turbidity at 600 nm for PSP-1 (a and b), PSP-2 (c and d), PSP-3 (e and f), PSP-4 (g and h), and PSP-5 (I and j). Approximate *C_sat_* values are indicated in green lines. Respective bright-field and confocal images of peptides in 50 mM tris buffer, 2.5 mM NaCl, pH 8.0, and in phase-separating concentrations of the peptide (indicated as dotted circles in the phase diagram) for PSP-1 (k and l), PSP-2 (m and n), PSP-3 (o and p), PSP-4 (q and r), and PSP-5 (s and t). (*n* = 3, scale bar = 20 µm). Images of Eppendorf tubes in phase separating conditions are shown as insets. Corresponding FRAP analysis for PSP-1 (u), PSP-2 (v), PSP-3 (w), PSP-4 (x), and PSP-5 (y). (*n* = 3, scale bar in FRAP insets = 2 µm).

### Capping of thiols prevents condensate formation

To uncover the precise role of cysteine thiols in condensate formation, we investigated the ability of peptides to phase separate when thiol groups are rendered incapable of forming disulfide bonds. To do so, iodoacetamide (IA) was used to cap thiols covalently in phase-separating conditions with non-phase-separating conditions as controls (Figure 3a)^37^. The amount of IA was also varied (0.5 molar equivalence or excess) such that the cysteines were capped partially (one of the two cysteines) or completely (both cysteines). In all cases, the peptides were first capped with IA in non-phase separating buffer conditions (high concentrations without salt) and then were diluted to non-LLPS (low concentrations, no salt) or LLPS conditions (> *C_sat_*; 2.5 mM NaCl). The samples were then investigated by matrix-assisted laser desorption ionization-time of flight (MALDI-ToF) mass spectrometry and confocal fluorescence microscopy using 1% of FITC-tagged peptides in the samples. PSP-2 and PSP-3, which contain two cysteine residues under non-phase-separating conditions, showed heterogenous mixtures when capped partially with IA, as confirmed by MALDI-ToF spectra (first panel; Figures 3b and c). In excess conditions, PSP-2 displayed both singly and doubly capped peptides, while PSP-3 showed only a doubly capped peptide (second panel; Figures 3b and c). Images of the samples in a confocal microscope showed no droplets in these conditions as expected (insets in the top two panels; Figures 3b and c). In LLPS conditions, when partially capped, mass spectra of PSP-2 showed the presence of uncapped and singly capped peptides (third panel; Figure 2b).

**Figure 3:**
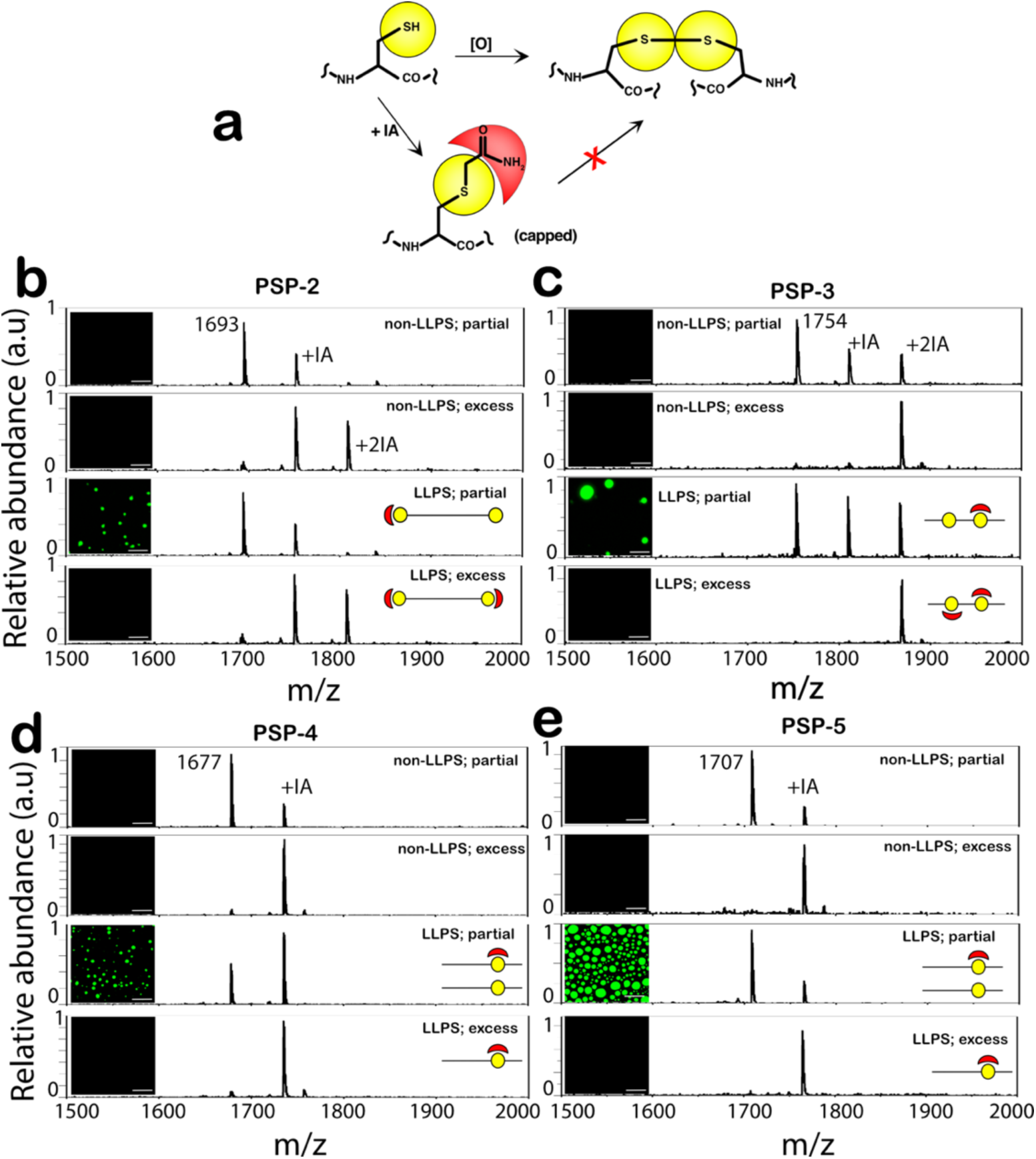
Preven-ng thiol oxida-on to disulfide bond inhibits condensate forma-on. (a) The schematic of the reaction involving Iodoacetamide (IA) capping of thiols. (b-e) The peptides PSP-2, PSP-3, and PSP-4 at 9mM and PSP-5 at 30 mM concentrations were incubated for two hours in water at room temperature (non-LLPS conditions) either with 0.5 molar excess of IA for partial capping or with a two-fold molar excess of IA for complete capping. The peptide stocks were then diluted to 3.0-3.5 mM either in LLPS conditions (20 mM Tris, 2.5 M NaCl, pH 8.0) or non-LLPS conditions (20 mM Tris, pH 8.0) followed by the analysis by MALDI-ToF mass spectrometry (MS) and confocal microscopy. MS spectra with partial and full-capping in non-LLPS conditions (top and second panels, respectively) and LLPS conditions (third and bottom panels, respectively) for (b) PSP-2, (c) PSP-3, (d) PSP-4, and (e) PSP-5. Insets in the panels show corresponding confocal microscopy images (scale bar = 20μm).

Under these conditions, the peptides showed condensate formation (inset in the third panel; Figure 2b). However, under fully capped and phase-separating conditions containing singly and doubly capped populations, the peptide failed to form condensates (fourth panel; Figure 3b). Similarly, under partial capping conditions, PSP-3 showed the presence of a heterogeneous mixture of uncapped, singly capped, and doubly capped peptides, which formed condensates (third panel; Figure 3c). However, when fully capped with excess IA, PSP-3 failed to phase separate (fourth panel; Figure 3c), which confirms that some populations of disulfide-bonded peptides are required for LLPS. PSP-4 and PSP-5, which contain single cysteines, showed no phase separation under non-LLPS conditions regardless of partial or full capping (first and second panels; Figures 3d and e). When partially capped under LLPS conditions, both peptides showed condensate formation (third panels; Figures 3d and e), but when fully capped, they failed to form droplets (fourth panels; Figures 3d and e). From these data, one can infer that thiol oxidation to disulfide crosslinks is key to driving the observed LLPS and forming the condensates.

### Condensates with disulfide crosslinks are reversible under redox gradients

If disulfide bonds are crucial for condensate formation, we questioned whether the condensates can reversibly form and dissolve under redox flux. To investigate this, peptide condensates were generated in respective phase separation conditions (3.5 mM peptides, 2.5 mM NaCl, and pH 8.0 for PSP-2 and PSP-3; 3.5 mM peptide, 2.5 mM NaCl, and pH 8.0 for PSP-4, and 18 mM peptide, 3.0 mM NaCl, 1% H_2_O_2_, and pH 8.0 for PSP-5). All peptides showed turbidity due to droplet formation on confocal microscopy (Figure 4, oxidized panels). The condensates were then titrated with increasing concentrations of dithiothreitol (DTT) as a reducing agent. Even low concentrations of DTT significantly reduced the number of droplets of PSP-2 condensates formed in oxidizing conditions (Figure 4a). Increasing the concentration of DTT to 3 and 5 mM completely dissolved the condensates (Figure 4a), further affirming the significance of disulfide bonds in condensate formation. To see whether dissolved condensates can be assembled by changing the reducing environment to an oxidized one, 2% hydrogen peroxide was added to the same reaction mixture and imaged under a confocal microscope. PSP-2 droplets reappeared in the solution (2% H_2_O_2_; Figure 4a), with the solution turning turbid (inset within Figure 4a; 2% H_2_O_2_). FRAP analysis of the droplets before adding DTT and after reoxidation showed nearly identical recovery kinetics, suggesting that the reformed droplets had similar viscosity and liquid-like character (Figure 4b). Nearly identical behavior was also observed with the peptides PSP-3, PSP-4, and PSP-5 peptides (Figures 4c-h). These results unequivocally establish that disulfide bonds are critical for the formation of peptide self-coacervates, and the bonds can be controlled by redox gradients.

**Figure 4:**
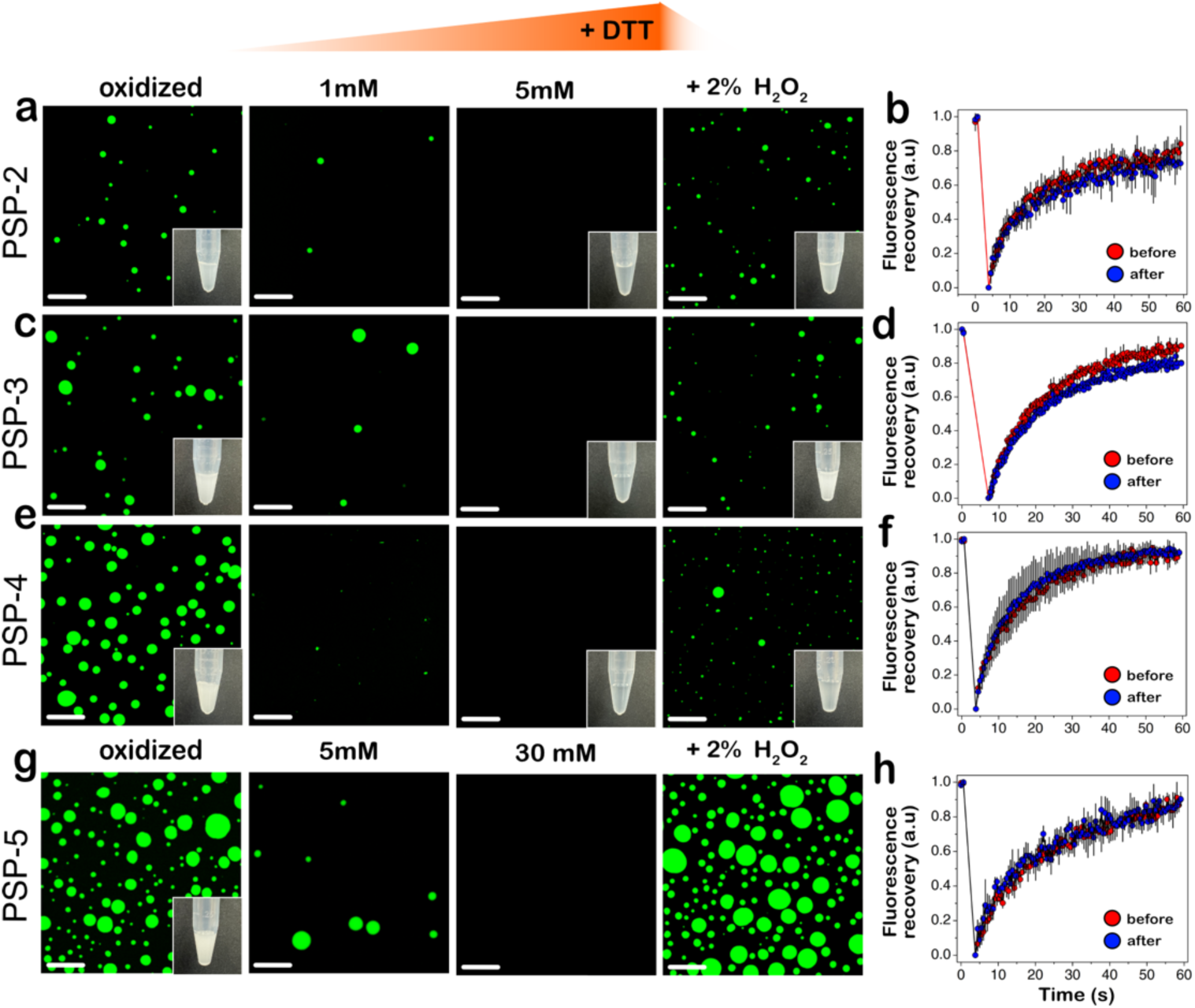
Peptide self-coacervates are redox-sensitive and reversible. Confocal microscopy images of FITC-labelled peptide self-coacervates in their respective LLPS conditions were titrated with DTT to reduce peptides and then reoxidized with hydrogen peroxide, followed by subsequent FRAP analysis. Confocal imaging with turbidity insets and FRAP analysis of the reduced peptide (before) and after oxidation (after) for PSP-2 (a and b), PSP-3 (c and d), PSP-4 (e and f), and PSP-5 (g and h). Insets show the images of Eppendorf tubes with samples in respective buffer conditions. (*n*=3, Scale bar = 20µm).

### Condensate fluidity and viscosity subtly vary with the positional variance of disulfide crosslinks

To fully understand whether the peptide condensates form system-spanning networks, interactions that are key for viscoelasticity ^25,45^, we monitored and measured the fluidity, morphology, and viscosity changes of the peptide condensates for 10 days at room temperature. Immediately after formation, the droplets of PSP-1 showed somewhat mitigated FRAP recovery, suggesting that they are highly percolation-prone, as mentioned above (Figure 2u). After one day, PSP-1 droplets showed significant coalescence, which further increased by three days (Figure 5a). Percolation was confirmed by the formation of a gel (Figure 5b). PSP-1 also showed percolation above 250 mM concentration without salt immediately after incubation (Figure 5c). This behavior is called percolation without phase separation^46^.

**Figure 5:**
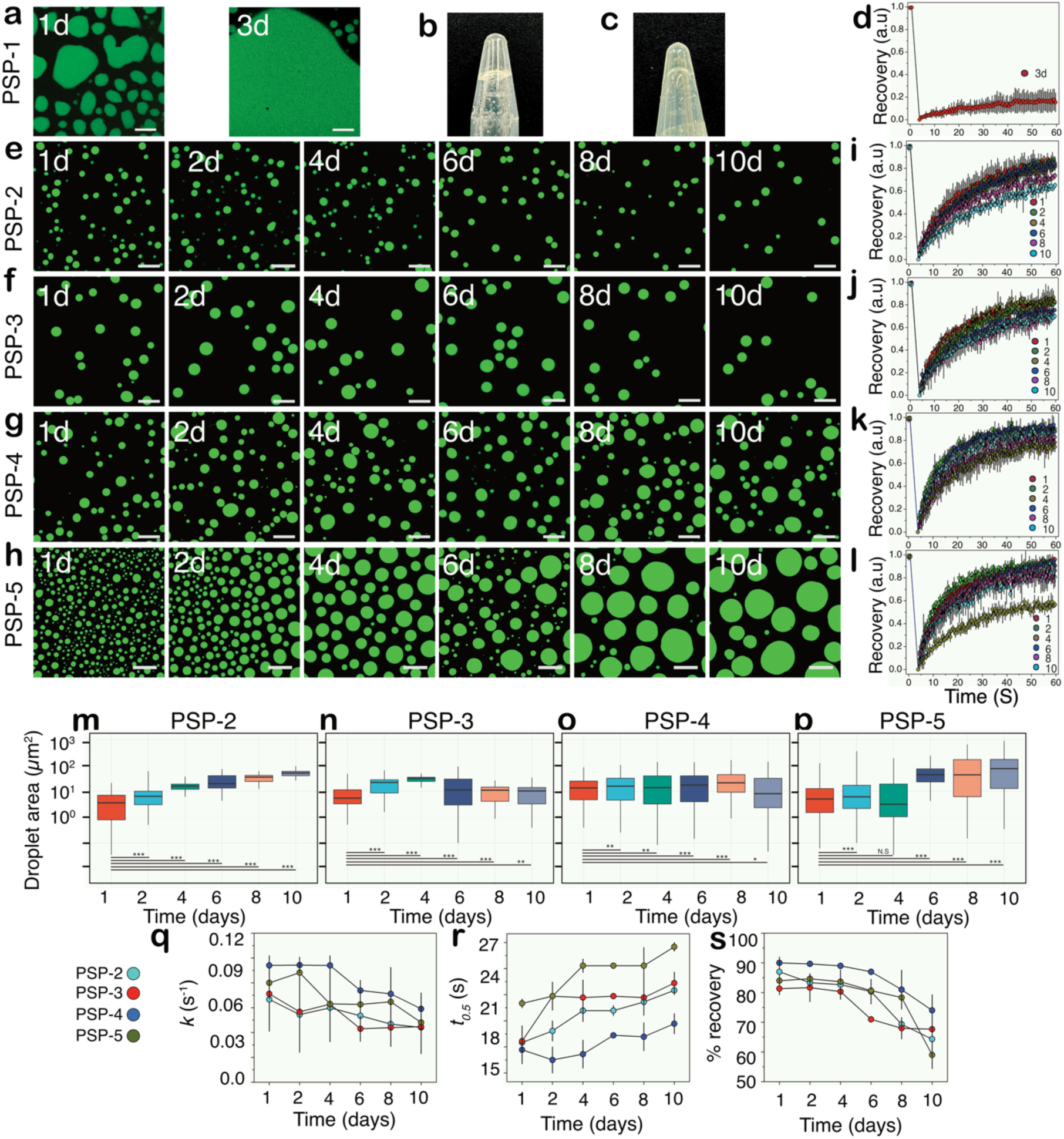
Viscoelastic changes of peptide self-coacervates over long incubation times. (a) Confocal microscopy images of PSP-1 self-coacervates (80 mM in 50 mM Tris, 3.5 mM NaCl, pH 8.0) after 1 and 3 days of incubation at room temperature. (b) An inverted Eppendorf tube containing the sample of PSP-1 after 3 days showing gelation. (c) An inverted Eppendorf tube image containing a 250 mM sample of PSP-1 in the absence of salt showing percolation without phase separation. (d-g) Confocal images of peptide coacervates (in respective LLPS conditions) at room temperature over 10-day incubation, along with their respective FRAP analysis for PSP-2 (d and h), PSP-3 (e and i), PSP-4 (f and j), and PSP-5 (g and k). Droplet size as a function of time for PSP-2 (l), PSP3 (m), PSP-4 (n), and PSP-5 (o). (\*\*\**p* <0.001, \*\**p* < 0.01, \**p* <0.05, N.S. = non-significant). First order rate constant (*k*) derived from the FRAP data (q), *t_0.5_* values (r), and percentage FRAP recovery (o). (*n* = 3, Scale bar of images=20 µm, FRAP insets = 2 µm).

Nevertheless, after three days of incubation, the samples showed negligible FRAP recovery (Figure 5d), suggesting that the droplets had crossed the percolation threshold to form system-spanning networks as expected for a model condensate. PSP-2 formed droplets with a surface area of ∼1-5 μm^2^ on the first day of incubation and, over 10 days, showed an average increase in the surface area to 30-40 μm^2^ (Figure 5e and m). This increase can be attributed mainly to the coalescence of the droplets, but importantly, the droplet morphology was maintained during the incubation period. The condensate viscosity was measured by FRAP analysis, which showed a rather consistent fluorescence recovery for 10 days, suggesting a liquid-like behavior (Figure 5i). During this time, the parameters, such as the rate constant for first-order exponential recovery (*k*), remained at ∼0.06 s^-1^ while the time taken to reach half recovery (*t_0.5_*) showed an insignificant increase from 18 to 21 s (Figures 5q and r). This data was consistent with the percentage recovery that decreased from 87% to 65% in the 10-day period (Figure 5s). PSP-3 and PSP-4 showed a droplet surface area of ∼10 μm^2^ that remained unchanged for 10 days (Figures 5f, g, n, and o). The viscosity of the condensates showed minimal change by FRAP (Figures 5j and k). While the rate constant, *k*, for PSP-3 decreased from 0.07 to 0.04 s^-1^, and a decrease from 0.09 to 0.06 s^-1^ was observed for PSP-4 (Figure 5q).

Similarly, the *t_0.5_* values averaged between 21 and 18 s for PSP-3 and PSP-4, respectively (Figure 5r). The percentage of FRAP recovery decreased from 80% to 70% for PSP-3 and decreased from 90% to 75% for PSP-4 within the 10-day incubation period (Figure 5s). PSP-5 condensates, on the other hand, showed some distinct differences from the others. It showed the largest increase in the droplet surface area from 5 to 100 μm^2^ in 10 days (Figures 5h and p), indicating that these condensates have greater fluidity based on coalescence alone. However, while *k* and *t_0.5_* did not show significant changes that averaged at 0.06 s^-1^ and 24 s, respectively, the percentage recovery decreased from 84% to 59% (Figures 5q-s). Interestingly, the largest decrease in FRAP recovery was observed between 8 and 10 days, suggesting that the PSP-5 condensates may have started to form percolation networks or, in other words, become more viscoelastic (Figure 5s). These data reveal that the positional and compositional variance of cysteines in the peptides subtly alter the fluidity and gelation of the condensates but largely remain unchanged. In other words, the disulfide bond crosslinks inhibit ‘gelation’, and help maintain the fluidity of the condensates.

### Peptides show conformational differences in phase-separating and homogenous buffer conditions

To see whether the peptides adopt different conformations within condensates and in bulk solution, we investigated by ^1^H nuclear magnetic resonance (NMR) spectroscopy and far-UV circular dichroism (CD). PSP-1, the control peptide in non-LLPS conditions, showed the expected aromatic chemical shifts at 7.0 and 7.3 ppm from the tyrosine residues and backbone amide protons between 7.6 and 8.7 ppm (black; Figure 6a). Chemical shifts of other protons were evident between 3.0 and 4.2 (Figure 6b) and 1.6 and 2.1 ppm (Figure 6c). PSP-2 showed significant changes in the chemical shifts in all protons in LLPS and non-LLPS conditions (red; Figure 6 a-c). Similarly, the PSP-3 (orange) and PSP-4 (green) peptides also showed changes in ^1^H chemical shifts between LLPS and non-LLPS conditions (Figure 6 a-c).

**Figure 6:**
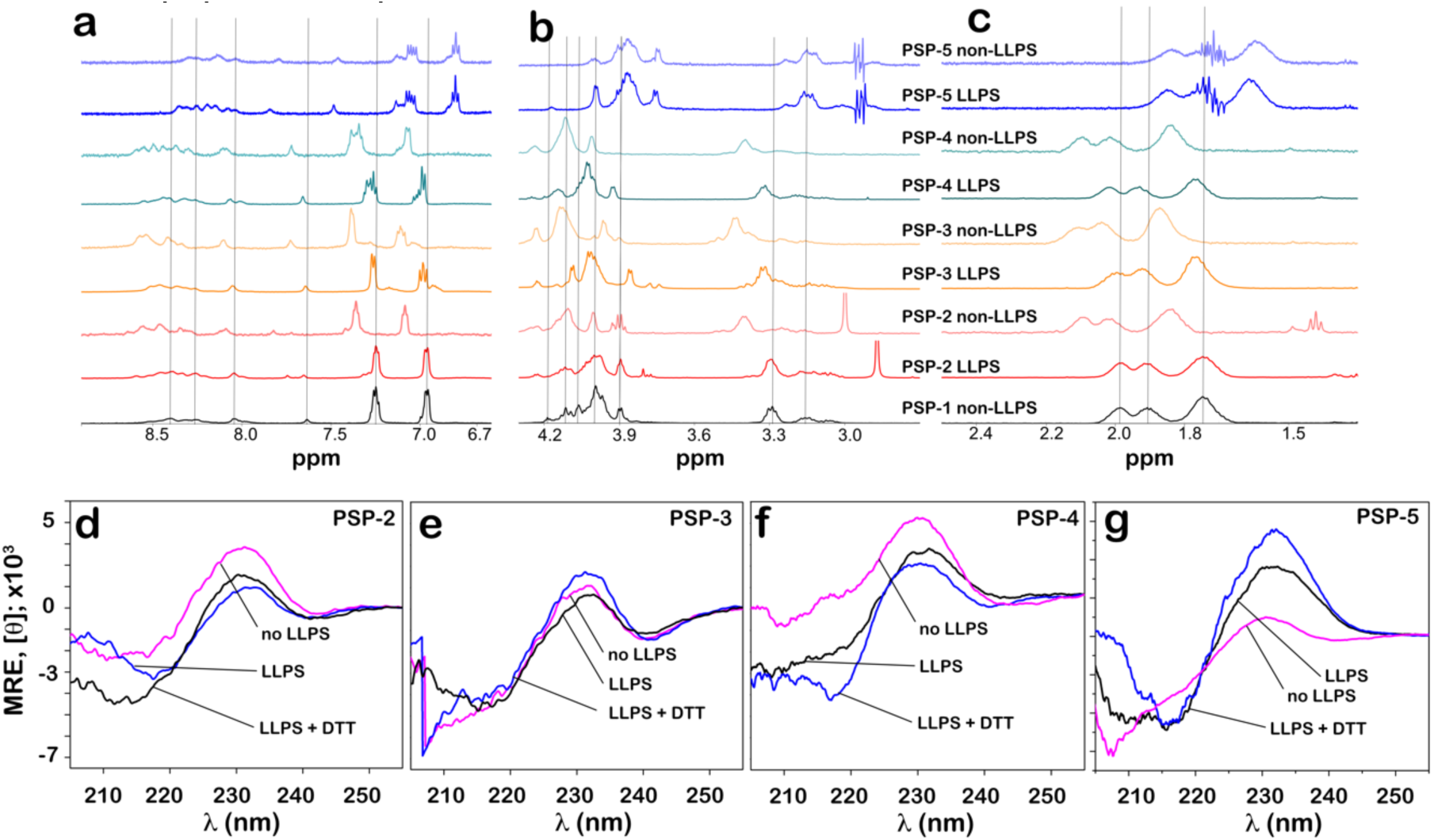
Conforma-ons of the pep-de show differences in bulk and condensed phases. Structural analysis of the peptides. (a-c) 1D ^1^H NMR spectra of the peptides in respective LLPS and non-LLPS conditions in 50 mM phosphate, 2.5 mM NaCl, pH 8.0 at room temperature as follows: PSP-1 non-LLPS = 1 mM, 2.5 mM NaCl; PSP-2 non-LLPS = 1 mM, PSP-2 LLPS = 3.5 mM; PSP-3 non-LLPS = 1.0 mM; PSP-3 LLPS = 3.5 mM; PSP-4 non-LLPS = 1 mM; PSP-4 LLPS = 3.5 mM; PSP-5 non-LLPS = 3.5 mM; PSP-5 LLPS = 18 mM. The spectra show the aromatic and backbone amide region (a) and other regions (b and c). Vertical lines indicate chemical shifts that vary significantly between LLPS and non-LLPS conditions. (d) CD spectra for PSP-2 at non-LLPS conditions (3.5 mM PSP-2 at 0.5M NaCl, pH-8.0), at LLPS conditions (3.5 mM PSP-2 at 2.5 M NaCl, pH-8.0) and at reducing conditions (with 4mM DTT). (e) CD spectra for PSP-3 at non-LLPS conditions (3.5 mM PSP-3 at 0.5M NaCl, pH-8.0), at LLPS conditions (3.5 mM PSP-2 at 2.5 M NaCl, pH-8.0) and at reducing conditions (with 4mM DTT). (f) CD spectra for PSP-4 at non-LLPS conditions (3.5 mM PSP-4 at 0.5M NaCl, pH-8.0), at LLPS conditions (3.5 mM PSP-2 at 2.5 M NaCl, pH-8.0) and at reducing conditions (4mM DTT). (g) CD spectra for PSP-5 at non-LLPS conditions (4 mM PSP-5 at 3.5M NaCl, pH-8.0), at LLPS conditions (18 mM PSP-5 at 2.5 M NaCl, pH-8.0), and at reducing conditions (60mM DTT).

However, PSP-5 showed more differences in the amide region (7.8 – 8.6 ppm) between LLPS and non-LLPS conditions than in other regions (blue; Figure 6 a-c). Together, the data indicates that the peptides adopted different conformations within the condensate and in dilute phases. The NMR results were also supported by far-UV CD, which provided time-averaged conformation of the peptide. All peptides showed a positive maximum at 232 nm, indicating a turn conformation (Figure 6 d-g). In addition, PSP-2 showed a negative minimum at 218 nm, indicating a β-sheet conformation in LLPS condition that was absent in non-LLPS conditions (pink and blue; Figure 6d). Reduction of the LLPS sample with DTT led to the loss of β-sheet, but the turn conformation remained (black; Figure 6d). PSP-3 did not show appreciable deviation from the turn conformation in LLPS, non-LLPS, and reducing conditions (Figure 6e). PSP-4 only demonstrated a change from turn to β-sheet under reducing conditions (Figure 6f). PSP-5 showed only a turn conformation in non-LLPS conditions but a β-sheet conformation alongside turns in LLPS conditions (Figure 6g). This conformation change was reflected in the backbone amide shifts in NMR. Under reducing conditions, the peptide showed partially disordered conformation (black; Figure 6g). Together, the data indicate that peptides adopt distinct conformations within and outside the condensates.

### Complex-coacervation with RNA markedly decreases the *C_sat_* values and renders the condensates insensitive to redox flux

In cells, BCs, such as stress granules, p-bodies, and others, are formed primarily via complex-coacervation involving several proteins and RNA molecules^47,48^. To assess the propensity of the designed peptides to undergo complex-coacervation with RNA, we sought to establish a phase diagram by scanning LLPS conditions across a concentration landscape of peptides and poly-A RNA under both an oxidizing and reducing environment (Figure 7). Surprisingly, PSP-1, the control peptide lacking cysteines, showed condensates in the presence of RNA with a low *C_sat_* of 75 μM in 250 μg/mL of RNA under oxidizing conditions (-DTT; Figure 7a), an order of magnitude reduction from the observed self-coacervation *C_sat_* of 80mM observed in the absence of RNA. As expected, under reducing conditions, the peptide did not show any change in the phase boundary since PSP-1 is devoid of cysteine residues (+DTT; Figure 7a). PSP-2-RNA condensates also showed a *C_sat_* value between 50 and 75 μM in 250 μg/mL of RNA in oxidizing conditions, which is almost an order of magnitude lower than that for self-coacervation (2.0 mM) (-DTT; Figure 7b). More interestingly, the phase boundary did not change in fully reducing conditions, suggesting that the condensates are insensitive to disulfide bond crosslinks (+DTT; Figure 7b). An identical phase boundary and redox insensitivity were also observed for PSP-3, PSP-4, and PSP-5 with respective *C_sat_* values between 50 and 75 μM in 250 μg/mL of RNA in oxidizing conditions (- and +DTT; Figures 7c-e). In oxidizing conditions, PSP-1 samples, in the condition indicated in a dotted circle in Figure 7a (250 μM with 200 μg/mL RNA), showed turbidity and numerous small droplets in the range of 0.5-3 μm diameter (Figure 7f). Nearly identical behavior was observed with peptides PSP-2, PSP-3, PSP-4, and PSP-5 under similar conditions (Figures 7g-j). PSP-1 showed somewhat attenuated FRAP recovery (∼ 60%), potentially revealing viscosity within the condensates (Figure 7k). Droplets of RNA coacervates with other peptides showed FRAP recoveries between 60 and 80% (Figures 7l-o). The data revealed that the peptides efficiently undergo complex coacervation with RNA. Furthermore, significantly smaller droplets were observed with complex-coacervates than self-coacervates of the peptides (Figure S3). Among the peptides, PSP-4 and PSP-5 showed the largest, and PSP-2 and PSP-3 showed more modest droplet size differences (Figure S3). Surprisingly, complex-coacervation with RNA renders the condensates insensitive to redox flux, likely due to multivalent electrostatic interactions overcompensating the effects of disulfide crosslinks. This point is discussed in further detail below.

**Figure 7:**
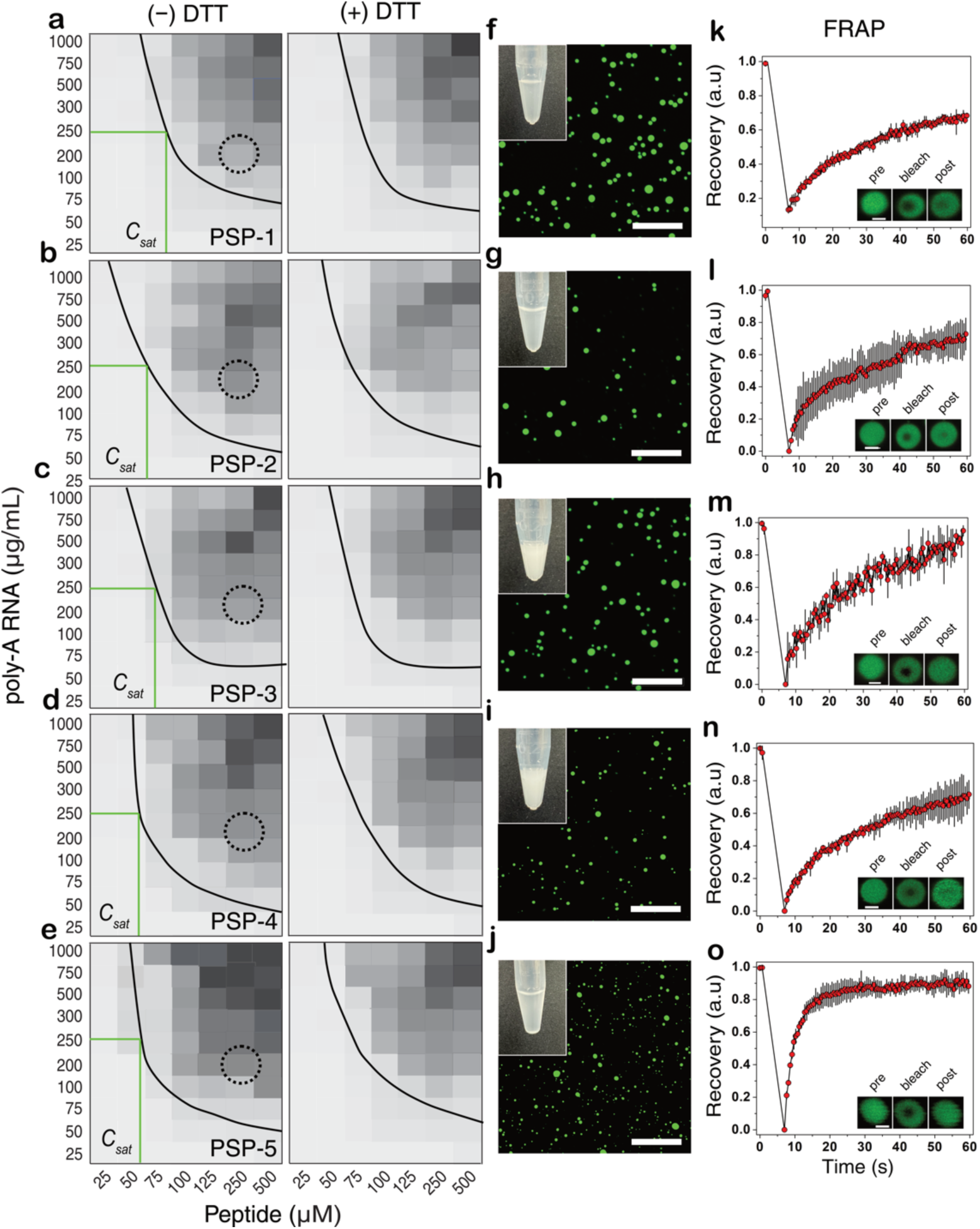
Peptide-RNA complex-coacervates are redox-insensitive. (a-e) Phase boundaries of poly-A RNA and PSP-1-5 complex-coacervates, respectively, in oxidizing (-DTT) and reducing (+DTT) conditions in 50 mM Tris pH 7.4. (f-j) Confocal microscopy images of FITC labeled peptides in concentration conditions indicated with dotted circles in panels (a-e; 250 µM peptide, 200 µg/mL poly-A RNA in 50 mM Tris, pH 7.4). Insets show the pictures of Eppendorf tubes with the corresponding samples. (k-o) Results from fluorescence recovery after photobleaching (FRAP) performed on the droplets shown in panels (f-j). (n=3, Scale bar of images = 10 µm, FRAP insets = 2 µm).

### Disulfide cross-linked condensates can be compartments for molecular cargo in peptide self-coacervates

Given the tunability of the condensate properties based on disulfide crosslinks, we questioned whether the condensates could host molecular cargo limited by their pore size and electrostatic charges on the cargo. To do so, we first evaluated the pore size of the condensates formed by peptides containing different disulfide bond crosslinks.

Using fluorescently labeled dextran with varying sizes, the pore sizes were measured by their ability to partition^45^ (Figure S4. Based on this analysis, we determined that all peptide condensates are porous with sizes larger than 100 nm (Figure S4). To test whether partitioning is limited by molecular charge, fluorescein, and tetramethylrhodamine methyl ester (TMR-OMe) dyes, which have similar sizes but with negative and positive charges at pH 8.0, respectively, were used. We measured the “encapsulation efficiency (EE)” by a colorimetric assay schematically shown in Figure 8a (see Methods). Fluorescein and TMR-OMe were partitioned within the droplets of PSP-2, PSP-3, PSP-4, and PSP-5 and visualized on a confocal microscope (Figure 8b). UV-visible spectrometry was utilized to quantify the amount of dye in the supernatant (and, therefore, by deduction of the amount within the droplets) (Figures 8c and 8d). The amount of dye in the dense phase was then calculated indirectly from the total amount. The dilute phase in all peptide condensates showed a significant presence of fluorescein (Figure 8c). In contrast, TMR-OME showed low absorbance (Figure 8d), indicating substantially more TMR-OMe was accommodated within the droplets than fluorescein. The calculated EE suggested that all peptides accommodated TMR-OMe at least three-fold better than fluorescein (Figure 8e). While PSP-2, PSP-3, and PSP-5 condensates accommodated ∼ 90-95% of TMR-OMe, PSP-4 accommodated only 80% (Figure 8e). In contrast, all peptides accommodated only 18-35% of fluorescein (Figure 8e). Since all peptides are highly porous (> 100 nm; Figure S4), partitioning is not limited by the size of the two dyes but by the molecular charge on the cargo. The peptide prefers positively charged TMR-OMe over negatively charged fluorescein. Fluorescein seems to be less preferred within the droplets, yet the peptides complex-coacervate with negatively charged RNA. This is likely due to the compatibility of positively charged TMR-OMe for cation - π interactions, especially at sub-stoichiometric quantities.

**Figure 8:**
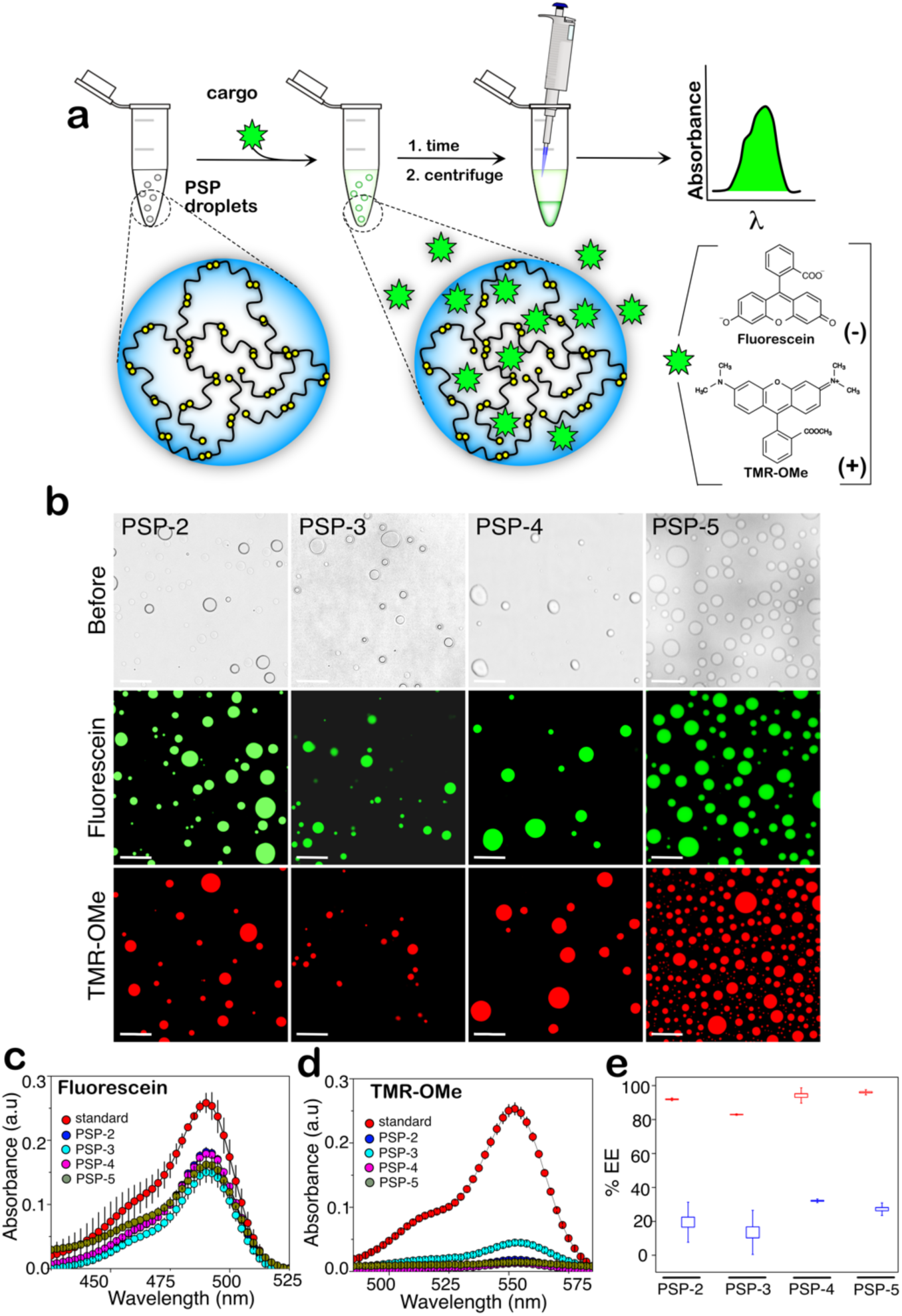
Pep-de condensates show different encapsulation efficiency for partitioning payloads. (a) Schematic of the assay performed to obtain encapsulation efficiency (EE) for fluorescein and tetramethyl rhodamine (TMR-OMe) payloads. (b) Bright-field images before adding dye (top) and confocal microscopy images of dye partitioning within PSP-2, PSP-3, PSP-4, and PSP-5 peptide condensates (bottom). The scale bar is 20 µm. (c and d) The average of at least three independent UV spectra of the supernatants was obtained after centrifugation (see (a)) after incubation with fluorescein and TMR-OMe. (d) The measured EE for fluorescein (blue) and TMR-OMe (red) dyes within the peptide droplets was calculated from the absorbance measurements in (c and d).

## CONCLUSIONS

The results presented here provide several novel conclusions on the role of cysteine disulfide bonds in the formation and viscoelasticity of biomolecular condensates, which has remained unknown thus far. First, disulfide bonds formed by the cysteines interspersed within a canonical sticker spacer framework of peptide promote condensate formation by decreasing the peptides’ saturation concentrations (*C_sat_*). Second, the redox chemistry of thiols undergoing disulfide bonds controls the reversibility of the condensates formed. The empirical observations presented here raise an important question: Are cysteines covalent stickers or auxiliary spacers in the stickers and spacers model of peptide condensates? The answer to this question is far from trivial. If disulfide bonds are stickers, the strong covalent bonds violate the prerequisite of multivalent weak interactions for condensate formation. On the other hand, if the cysteines are present within the spacers, they are likely to affect the effective solvation volume, which is also key for LLPS^16^. However, we believe that cysteines function neither as stickers nor spacers but as nodes for crosslinks that enhance intermolecular sticker-sticker interactions by decreasing the effective concentrations. They also help maintain liquid-like characteristics for a prolonged period of time. This can be reconciled from the four categories of peptides that can be classified I-IV based on their cysteine compositions (Figure 9). Classes I and II can form an extended network of covalently bonded disulfide crosslinks at high peptide concentrations in addition to some dimers in an oxidized state. Classes III and IV can exclusively form dimers. Therefore, class I and II peptides (PSP-2 and PSP-3), with their two cysteines on the termini and interior forming disulfide networks (Figure 9), render the stickers interactions to transition to predominantly intramolecular from intermolecular in the reduced state. The formation of the disulfide network significantly decreases the effective concentrations of the sticker interactions, which manifests in the reduction of *C_sat_*, as seen in Figure 2. As expected, the disulfide network-forming PSP-2 and PSP-3 peptides have lower *C_sat_* than the dimer-forming PSP-3 and PSP-4 peptides. Although disulfide crosslinks decrease *C_sat_* for peptide self-coacervates, under our oxidized and reduced experimental conditions, the phase boundary of complex-coacervates with RNA remains innocuous to redox flux (Figure 7). We conjecture that the sum of energies contributed by multivalent electrostatic interactions between RNA backbone and peptides exceeds the net decrease in effective concentrations induced by disulfide crosslinks. In addition, these interactions may also decrease cation-π interactions between lysines and tyrosines. Furthermore, the overall negative charge on the peptide-RNA condensates may also prevent DTT from interacting with the condensates (and S-S bonding) due to negative charge repulsion between the condensates and the DTT. However, several parameters, such as RNA concentrations, ionic strength, and pH, are likely to influence these dynamics, and our ongoing experimental analysis will decipher these aspects and will be reported later. Nevertheless, from the results presented here, one can unambiguously conclude that the peptide condensates are redox-sensitive to varying degrees, especially the self-coacervates.

**Figure 9:**
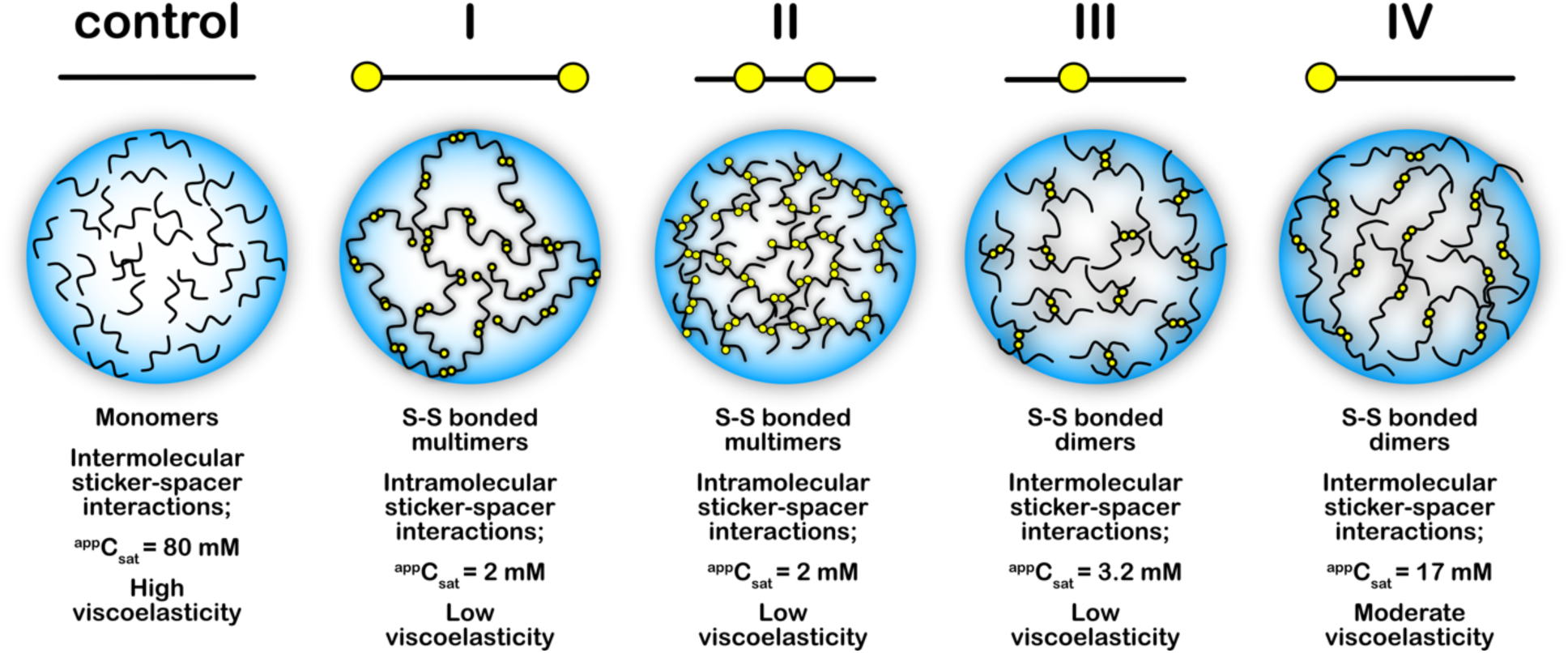
Overall conclusions

The data presented here provide insights into a fundamental understanding of cysteine’s role in the LLPS phenomenon, which may underlie mechanisms of several proteins in biology, especially those involved in pathologies involving LLPS. The results also showcase how cysteines can be incorporated into phase-separating model peptides and the design of customized, specific redox-tunable soft materials.

## AUTHOR INFORMATION

Conceptualization – VR; intellectual inputs – VR, MM, TDC, AKP; biophysical experiments, data interpretation – MM; peptide synthesis – PEJ, WSS, TDC; data processing – MM, LDL; NMR – DMD, TC, HW, AKP; manuscript writing and editing – VR, MM.

## ACKNOWLEDGMENTS

The authors thank the National Science Foundation (BMAT 2208349) for their financial support of this project. They also thank the National Center for Research Resources (5P20RR01647-11) and the National Institute of General Medical Sciences (8 P20GM103476-11) from the National Institutes of Health (NIH) for funding through INBRE to use their core facilities. Additionally, the authors acknowledge support from the National Science Foundation (Award number: 2319932) for mass spectrometry instrumentation utilized in this study. TDC acknowledges funding support from the NIH and the National Institute of Biomedical Imaging and Bioengineering (NIBIB R21EB033533). PEJ acknowledges fellowship support from the Mississippi Space Grant Consortium (MSSGC), funded by the National Aeronautics and Space Administration (NASA) Office of STEM Engagement.

## SUPPLEMENTARY FIGURES

**Figure S1.**
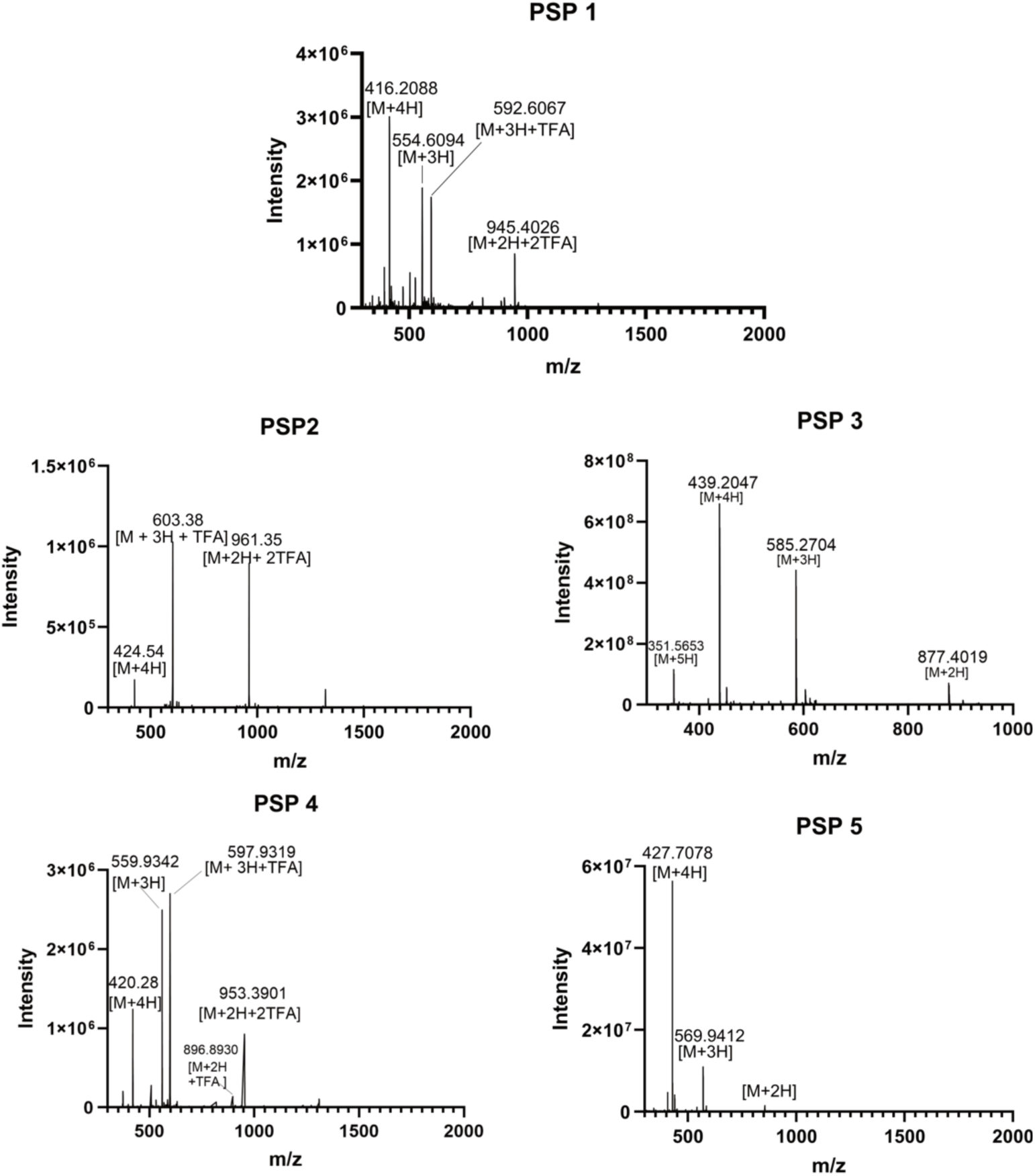
Confirmation of peptide synthesis and purity by ESI orbitrap mass spectrometry.

**Figure S2.**
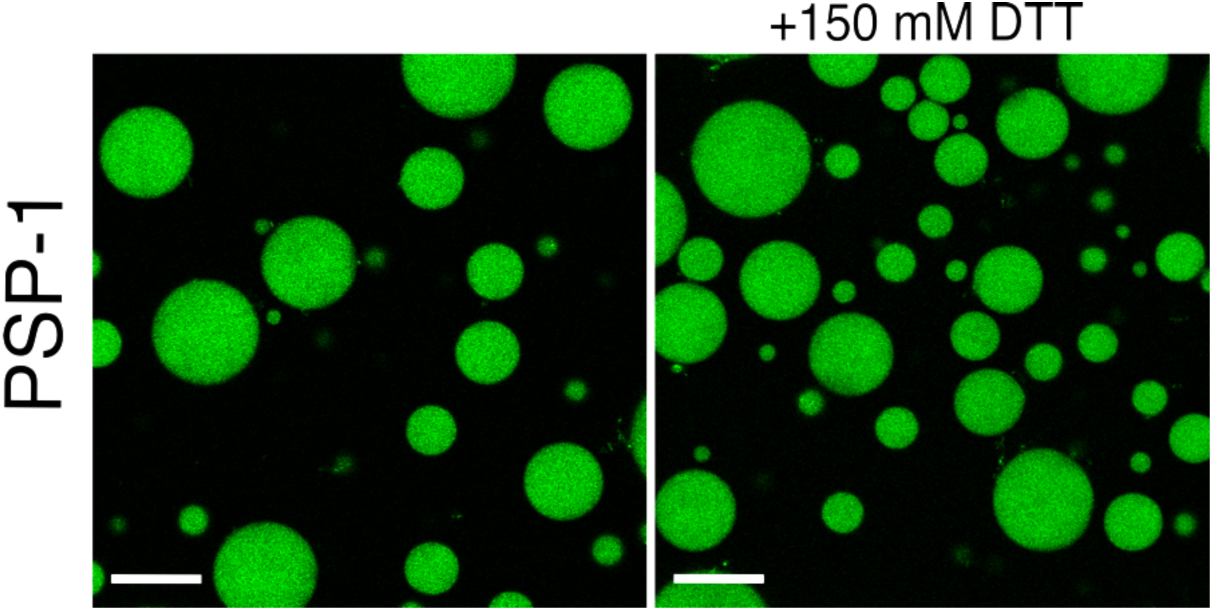
The addition of DTT does not change PSP-1 droplet morphology.

**Figure S3:**
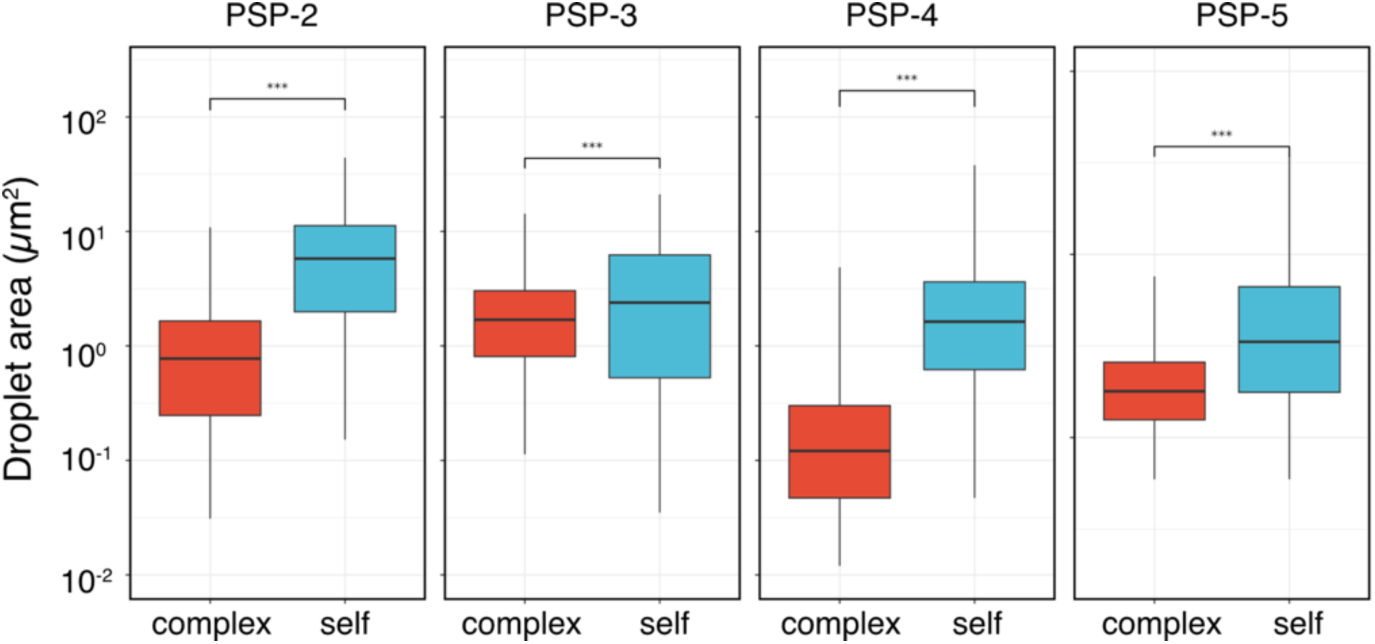
Self- and complex-coacervates show dis:nct differences in their droplet sizes. Box and whisker plots of peptide droplet surface area of complex- and self-coacervates obtained from confocal images. (*n*= 5; *** = p> 0.01).

**Figure S4.**
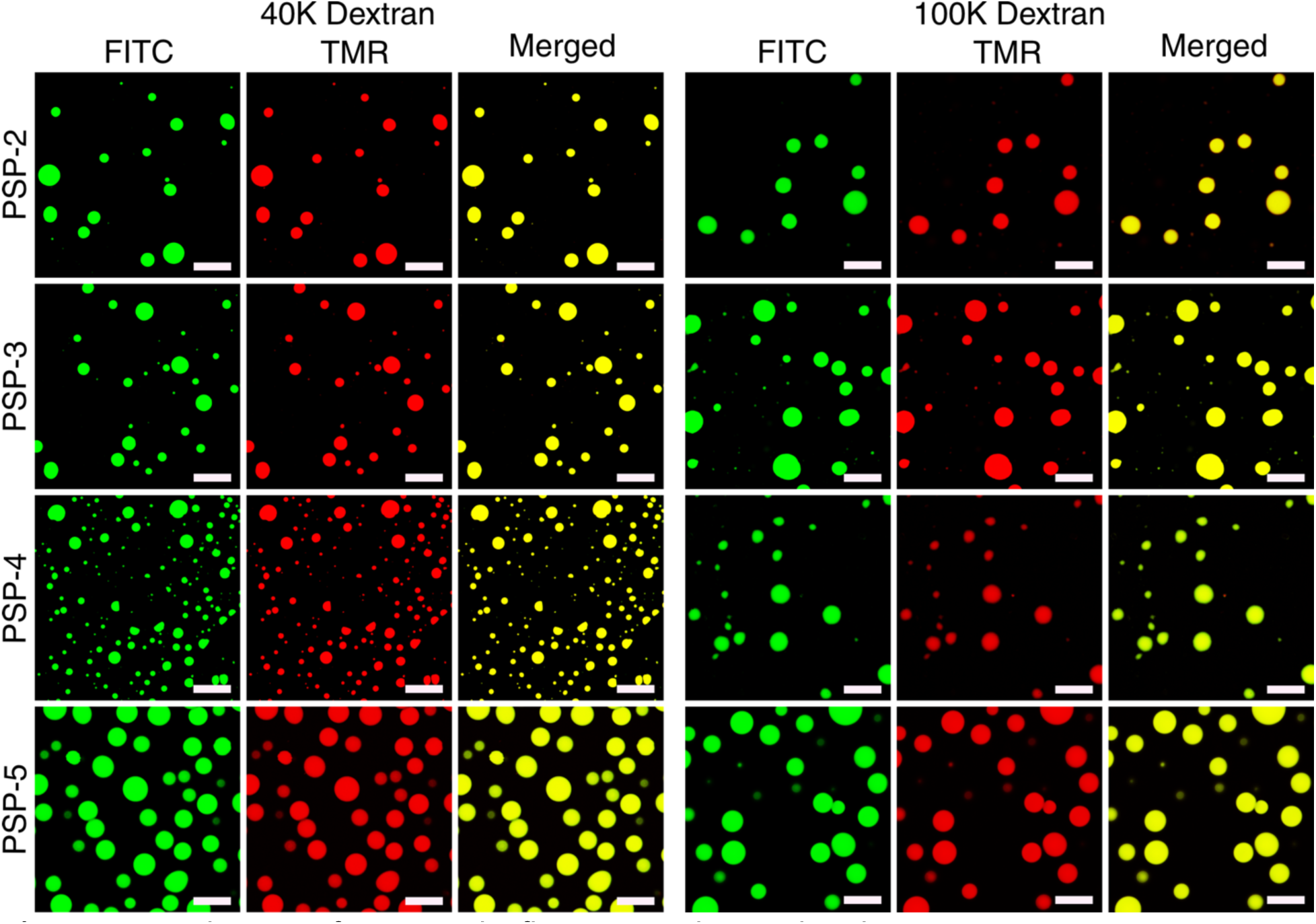
Evaluation of pore size by fluorescent dextran beads.

